# Early nuclear phenotypes and reactive transformation in human iPSC-derived astrocytes from ALS patients with *SOD1* mutations

**DOI:** 10.1101/2023.10.05.561079

**Authors:** Vincent Soubannier, Mathilde Chaineau, Lale Gursu, Sarah Lepine, David Kalaydjian, Ghazal Haghi, Guy Rouleau, Thomas M. Durcan, Stefano Stifani

## Abstract

Amyotrophic Lateral Sclerosis (ALS) is a neurodegenerative disease characterized by the progressive death of motor neurons (MNs). MN degeneration in ALS involves both cell-autonomous and non-cell autonomous mechanisms, with glial cells playing important roles in the latter. More specifically, astrocytes with mutations in the ALS-associated gene *Cu/Zn superoxide dismutase 1* (*SOD1*) promote MN death. The mechanisms by which *SOD1*-mutated astrocytes reduce MN survival are incompletely understood. In order to characterize the impact of *SOD1* mutations on astrocyte physiology, we generated astrocytes from human induced pluripotent stem cell (iPSC) derived from ALS patients carrying *SOD1* mutations, together with control isogenic iPSCs. We report that astrocytes harbouring *SOD1*(A4V) and *SOD1*(D90A) mutations exhibit molecular and morphological changes indicative of reactive astrogliosis when compared to matching isogenic astrocytes. We show further that a number of nuclear phenotypes precede, or coincide with, reactive transformation. These include increased nuclear oxidative stress and DNA damage, and accumulation of the SOD1 protein in the nucleus. These findings reveal early cell-autonomous phenotypes in *SOD1*-mutated astrocytes that may contribute to the acquisition of a reactive phenotype involved in alterations of astrocyte-MN communication in ALS.

## 1. Introduction

Amyotrophic lateral sclerosis (ALS) is an incurable motor neuron (MN) disease characterized by the progressive degeneration of MNs in the cerebral cortex, brain stem and spinal cord, resulting in gradual muscle paralysis and ultimately death by respiratory failure (Mejzini et al., 2019; Kim et al., 2020). The majority of ALS cases are sporadic, with less than 20% of cases inherited through families (familial ALS). Multiple deleterious variants in numerous genes have been associated with familial ALS over the years, starting with the discovery of the first genetic mutation to cause ALS, affecting the *Cu/Zn superoxide dismutase 1* (*SOD1*) gene (Rosen et al., 1993). Together with *SOD1*, *chromosome 9 open reading frame 72* (*C9orf72)*, *TAR DNA binding protein* (*TARDBP)*, and *FUS RNA binding protein (FUS)* are the most frequently mutated genes in ALS, accounting for approximately 70% of familial ALS cases (Mejzini et al., 2019; Kim et al., 2020; Brenner and Freischmidt, 2022). Although these genes play multiple functions in disease pathogenesis, many of which remain to be fully elucidated, protein misfolding and accumulation of toxic aggregates is a common feature of the most common familial ALS mutations (Calabrese et al., 2022; Tran and Lee, 2022; Arnold et al., 2023). Perturbations of numerous mechanisms contribute to MN death in ALS, including, but not limited to, oxidative stress, RNA metabolism and protein homeostasis, nucleocytoplasmic trafficking, dynamics of ribonucleoprotein bodies, mitochondrial functions, and autophagy (Balendra and Isaacs, 2018; Burk and Pasterkamp, 2019; Mejzini et al., 2019; Prasad et al., 2019).

Although traditionally defined as a disease that affects vulnerable MNs, there is increasing evidence that the convergence of damage within multiple cell types is crucial to MN loss in ALS. Similar to other neurodegenerative diseases, ALS is characterized by extensive neuroinflammation involving astrogliosis, activation of microglia, and infiltration of peripheral immune cells at sites of neuronal degeneration (Beers and Appel, 2019; Cipollina et al., 2020). Reactive astrogliosis is a hallmark of *post-mortem* tissue from ALS patients and ALS mouse models (Schiffer et al., 1996; Hall et al., 1998; Shibata et al., 2001; Johann et al., 2015). Several studies with cultured cells provide evidence suggesting that astrocytes harbouring ALS mutations can be toxic to MNs *in vitro* (Di Giorgio et al., 2007; Nagai et al., 2007; Haidet-Phillips et al., 2011). The contribution of astrocytes to MN pathology in ALS is complex, depending on the stage of disease progression. It is hypothesized that astrocyte reactive transformation initially occurs as a neuroprotective response during early stages of ALS. At least some activated astrocytes can then gradually become neuroinflammatory during disease progression, contributing to neuronal degeneration. The deleterious effects of astrocytes on MNs in ALS may result from loss of supportive functions and/or gain of toxic activities, such as secretion of neuroinflammatory molecules (Meyer et al., 2014; Varcianna et al., 2019; Guttenplan et al., 2020; Van Harten et al., 2021; Arredondo et al., 2022).

Approximately 15% of familial ALS case are caused by mutations in the *SOD1* gene, which encodes an abundant and broadly expressed protein that catalyzes dismutation of superoxide to hydrogen peroxide and molecular oxygen thereby protecting cells from reactive oxygen species toxicity. SOD1 is present in the cytosol, mitochondria, peroxisomes and nuclei (Okado-Matsumoto and Fridovich, 2001; Bunton-Stasyshyn et al., 2015; Xu et al., 2022). A large number of genetic variants of *SOD1* have been identified in ALS patients. *SOD1* mutations usually result in gain-of-function effects in which the mutated SOD1 proteins acquire new toxic functions thought to derive from misfolding and an increased predisposition to aggregation. Although the pathogenic mechanisms of aggregated SOD1 in MNs remain to be fully elucidated, they affect key cellular processes such as scavenging of free radicals, mitochondrial function, axonal transport, protein quality control, and mRNA splicing, to name a few (Bunton-Stasyshyn et al., 2015; Abati et al., 2020; Kim et al., 2020; Peggion et al. 2022).

In addition to affecting MN physiology, *SOD1* mutations have an impact on astrocyte biology and the cross-talk between astrocytes and MNs. More specifically, astrocytes harbouring *SOD1* mutations decrease MN survival both *in vivo* and *in vitro* (Di Giorgio et al., 2007; Nagai et al., 2007; Marchetto et al., 2008; Meyer et al., 2014). This effect is mediated by cell-to-cell signaling between astrocytes and MNs (Nagai et al., 2007; Fritz et al., 2013; Urban et al., 2023). Little information is available on how *SOD1* mutations drive intrinsic changes within astrocytes that affect their cross-talk with MNs. We report that the *SOD1*(A4V) and *SOD1*(D90A) mutations are associated with enhanced astrocyte reactivity in the absence of other cell types. Reactive astrogliosis is concomitant with increased DNA damage and accumulation of SOD1 in the nucleus. These changes are preceded by signs of increased nuclear oxidative stress. These findings reveal early nuclear phenotypes in *SOD1*-mutated astrocytes that may contribute to the acquisition of a reactive phenotype involved in mechanisms of MN degeneration in ALS.

## 2. Results

### 2.1. Generation of ventral spinal cord-like astrocytes from *SOD1*-mutated human iPSCs

Induced cells with molecular features of ventral spinal cord astrocytes were generated from human iPSC lines derived from ALS patients harbouring *SOD1*(A4V) or *SOD1*(D90A) mutations as described (Soubannier et al., 2022). The A4V mutation is the most frequent *SOD1* mutation in North America, while the D90A mutation is the most prevalent in Europe (Kim et al., 2020). Matching gene-edited iPSC lines were used as controls (hereafter termed Iso(A4V) and Iso(D90A) to indicate which specific mutations were corrected). Immunocytochemistry and RT-PCR confirmed the initial generation of neural progenitor cells (NPCs) with caudal and ventral neural tube properties, such as expression of the cervical marker *HOXA5* and ventral marker *NKX6.1* (Supplementary Figure 1 – Fig. S1). Upon exposure to pro-astrogenic culture conditions, these validated NPCs generated robust numbers of cells exhibiting a fibrous morphology and the co-expression of typical astrocyte markers, such as GFAP and S100B, as early as 30 days after the start of *in vitro* differentiation (DIV30). Comparable yields of GFAP^+^/S100B^+^ cells were observed when *SOD1*-mutated NPCs or their isogenic counterparts were used (Fig. 1A-D). Exposure of Iso(A4V) astrocytes to a combination of tumour necrosis factor-alpha (TNF-α), interleukin 1-alpha (IL-1α), and complement component 1, subcomponent q (C1q), previously shown to promote astrocyte reactivation (Liddelow et al., 2017), resulted in upregulation of several astrogliosis marker genes, such as *complement component 1, subcomponent s (C1s)*, *complement component 3* (*C3*), *S100 calcium binding protein A10* (*S100A10*), *pentraxin 3* (*PTX3*), *serpin family g member 1* (*SERPING1*), *c-c motif chemokine ligand 2 (CCL2*), and C-X-*c motif chemokine ligand 10* (*CXCL10*) (Fig. 1E). Consistently, a number of cytokines and chemokines were over-secreted by Iso(A4V) astrocytes treated with TNF-α, IL-1α, and C1q, compared to untreated cells (Fig. 1F). These observations show that iPSC-derived astrocytes display properties similar to physiological astrocytes.

**Figure 1.**
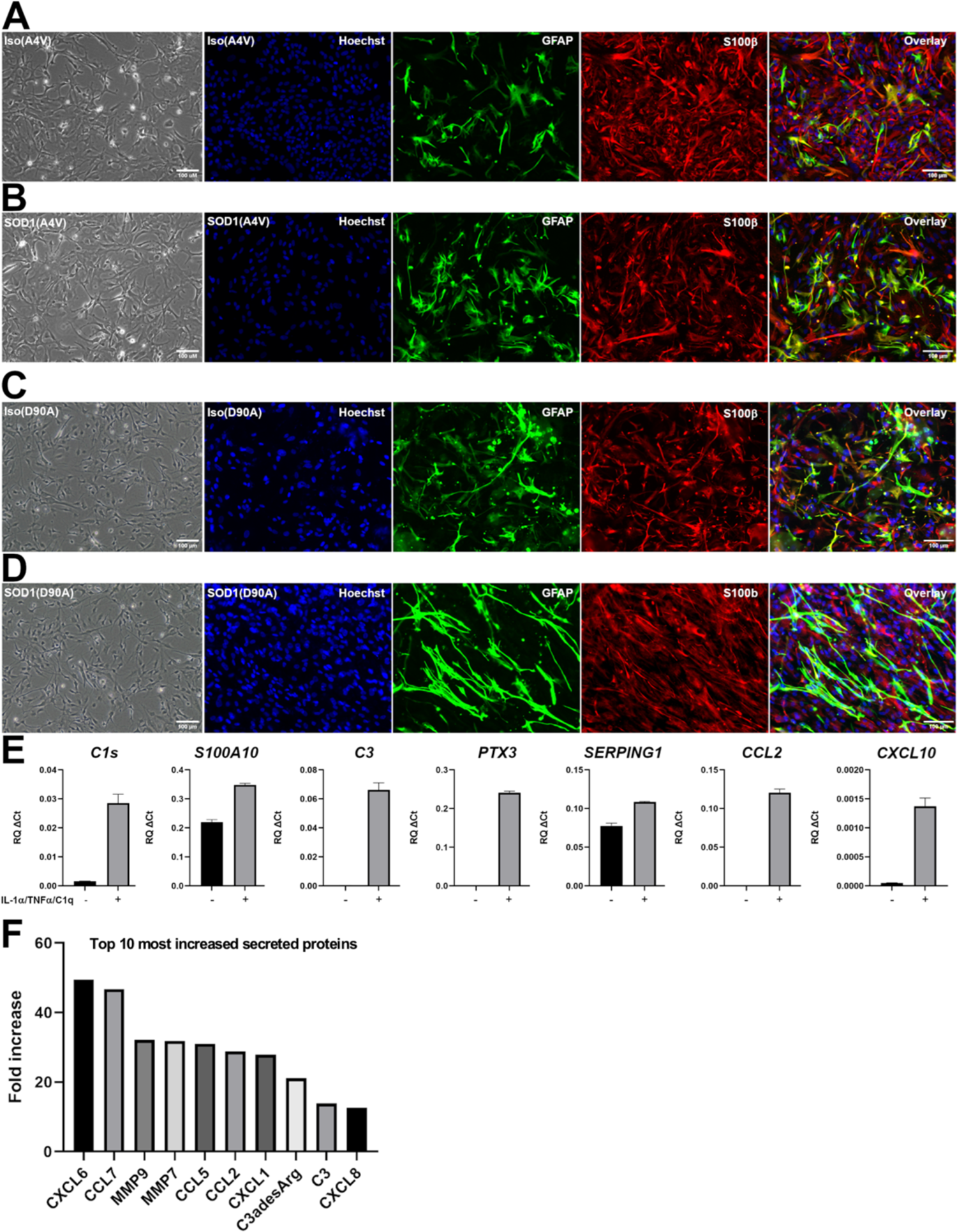
Characterization of iPSC-derived astrocytes. **A-D**) Representative images of phase contrast (left-hand panel in each row) and double-labeling immunofluorescence of GFAP (green) and S100B (red) expression in DIV60 astrocytes generated from iPSC lines with A4V or D90A mutations in *SOD1*, and their matching isogenic lines (termed Iso(A4V) and Iso(D90A) to indicate which mutations were corrected); Hoechst counterstaining (blue) is shown. The majority of induced cells express S100B, and a significant proportion of S100B-positive cells co-express GFAP at high level, while other S100B-positive cells co-express GFAP at lower levels. **E**) Real-time PCR analysis of the expression of the indicated reactive astrocyte markers in DIV60 Iso(A4V) astrocytes treated, or not, with IL-1α, TNF-α, and C1q. (**F**) List of the ten most increased cytokines and chemokines secreted in the medium by Iso(A4V) astrocytes following treatment with TNF-α, IL-1α, and C1q compared to treatment with vehicle. Differences are expressed as fold change.

### 2.2. *SOD1*-mutated astrocytes undergo reactive transformation

Previous studies have shown increased reactive transformation in astrocytes harbouring mutations in different familial ALS genes, including *SOD1*, *C9orf72*, *FUS*, and *valosin-containing protein* (*VCP)* (Birger et al., 2019; Taha et al., 2022; Stoklund Dittlau et al., 2023). Much remains to be learned about the cellular mechanisms underlying reactive astrogliosis in ALS, particularly cell-autonomous processes. In this context, we tested whether cultures of astrocytes harbouring the *SOD1*(A4V) mutation would undergo reactive transformation in the absence of extrinsic cues. Morphological comparison of *SOD1*(A4V) astrocytes and their isogenic counterparts using phalloidin staining to label cytoskeletal structures revealed no significant differences at DIV30 (Fig. 2A). Since reactive astrogliosis is usually characterized by a disassembly of F-actin stress fibers stained with phalloidin into a more disorganized G-actin network (Hansson, 2015; Tyzack et al., 2017), this observation suggested the lack of significant reactivation in *SOD1*(A4V) astrocytes at DIV30. To test this possibility further, we conducted quantitative RT-PCR studies to compare the expression levels of genes known to be up-regulated in reactive astrocytes (Liddelow et al., 2017). These studies showed a trend toward increased levels of *C1s* and *S100A10* in *SOD1*(A4V) astrocytes, but no significant difference in the expression of other reactive astrocyte phenotype markers, such as *C3*, *PTX3*, and *SERPING1* (Fig. 2B). These findings suggest that *SOD1*(A4V) astrocytes are not significantly more reactive than isogenic astrocytes after 30 days *in vitro*.

**Figure 2.**
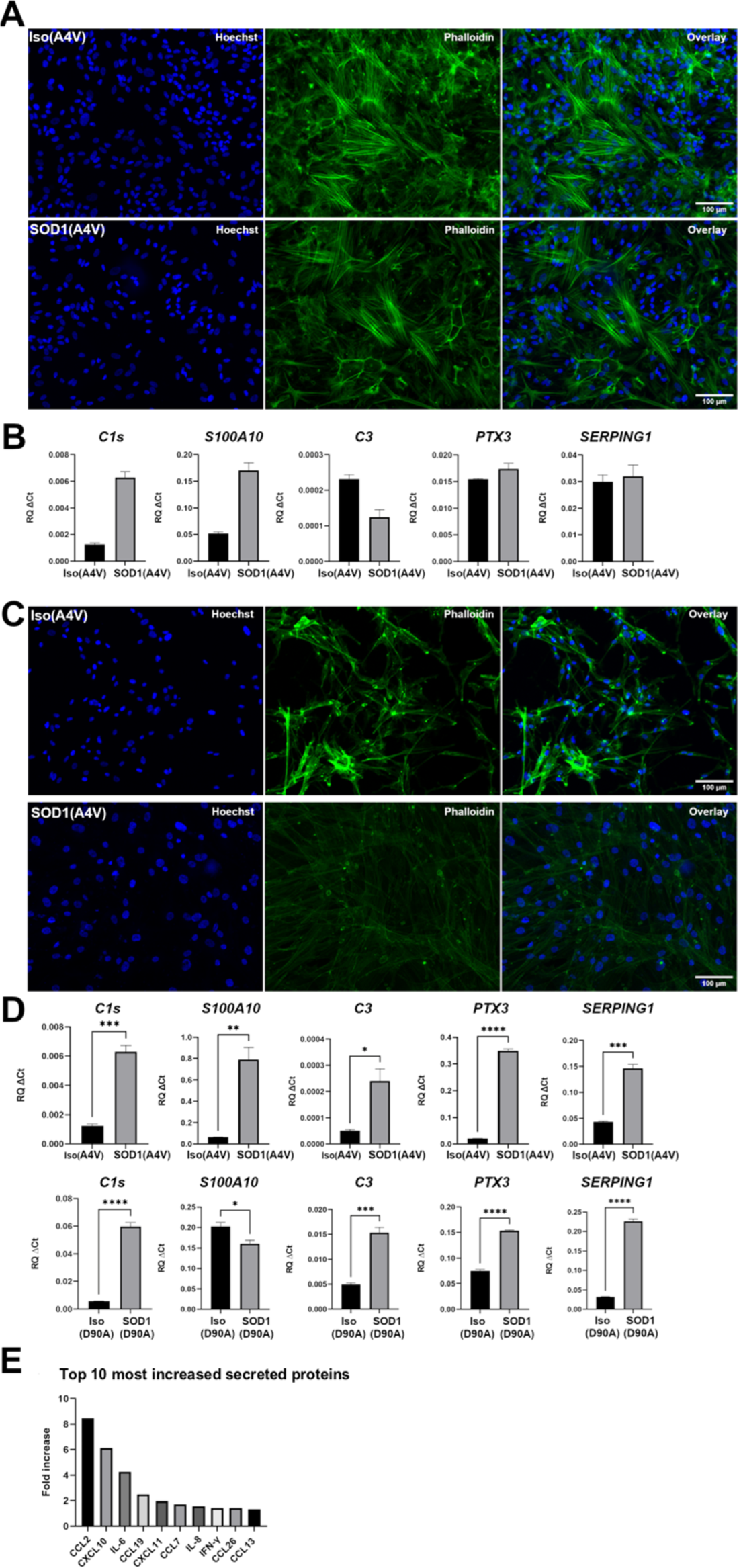
Reactive transformation of *SOD1*-mutated astrocytes. **A, B**) DIV30 astrocytes. A) Representative images of the actin cytoskeleton of *SOD1*(A4V) and matching isogenic astrocytes visualized by Alexa488-conjugated phalloidin staining (green); Hoechst counterstaining (blue) is shown. B) Real-time PCR analysis of the expression of the indicated reactive astrocyte markers in *SOD1*(A4V) and matching isogenic astrocytes. **C, D)** DIV60 astrocytes. C) Representative images of the actin cytoskeleton of *SOD1*(A4V) and matching isogenic astrocytes visualized by phalloidin staining (green); Hoechst counterstaining (blue) is shown. D) Real-time PCR analysis of the expression of the indicated reactive astrocyte markers in either *SOD1*(A4V) and isogenic astrocytes (top row) or *SOD1*(D90A) and isogenic astrocytes (bottom row). Statistical analyses were performed with Student’s t-test, graphs show mean ± *SEM*; \**p* < 0.05; \*\*\*\**p* < 0.0001; *n* = 3. **E)** List of the ten most increased cytokines and chemokines secreted in the medium by *SOD1*(A4V) astrocytes compared to Iso(A4V) astrocytes. Differences are expressed as fold change.

We next performed the same studies at DIV60, when induced astrocytes are more developmentally mature. Phalloidin staining showed that isogenic astrocytes continued to exhibit the presence of F-actin stress fibers typical of healthy cells, whereas *SOD1*(A4V) astrocytes displayed a disassembly of the stress fibers and the presence of actin networks characterized by ring-like structures, ruffles, and radial actin filaments, suggestive of reactive transformation (Fig. 2C). In agreement with this observation, *C1s*, *C3*, *S100A10*, *SERPING1*, and *PTX3* were all upregulated in *SOD1*(A4V)-mutated astrocytes at DIV60 (Fig. 2D). Moreover, *SOD1*(A4V) astrocytes exhibited increased secretion of several proteins associated with a reactive phenotype (Fig. 2E). Similar morphological and gene expression phenotypes were observed at DIV60 in astrocytes harbouring the *SOD1*(D90A) mutation, showing that these phenotypes were not unique to the A4V mutation in *SOD1* (Fig. 2C, D; Fig. S2). Together, these results provide evidence that astrocytes with *SOD1*(A4V) and *SOD1*(D90A) mutations undergo cell-autonomous reactive transformation by 60 days *in vitro* when compared to their isogenic counterparts.

### 2.3. Increased nuclear oxidative stress in *SOD1*-mutated astrocytes

To further characterize the phenotype of *SOD1*-mutated astrocytes, we first tested whether we could detect differences with isogenic astrocytes preceding detectable signs of reactive transformation. Based on previous studies showing increased oxidative stress in astrocytes carrying ALS mutations (Shibata et al., 2001; Birger et al., 2019; Appel et al., 2021), *SOD1*-mutated astrocyte cultures were tested for oxidative stress at DIV30 using a range of concentrations of the probe MitoSox, a dye readily oxidized by superoxide ions. In agreement with previous studies, we detected higher MitoSox levels in *SOD1*(A4V) astrocytes at a concentration of 1 μM (Fig. S3): this experimental condition is expected to detect mainly mitochondrial superoxide, a type of reactive oxygen species (Roelofs et al., 2015). Increasing the concentration of MitoSox to 5 μM, a dose at which this probe redistributes to the nucleus (Roelofs et al., 2015), revealed statistically significantly higher levels of nuclear MitoSox intensities in both *SOD1*(A4V) and *SOD1*(D90A) astrocytes, compared to the corresponding isogenic astrocytes (Fig. 3A, B). This finding suggests that astrocytes harbouring these ALS-associated *SOD1* mutations have increased generation, or impaired clearance, of superoxide ions within the nucleus as early as 30 days after the start of *in vitro* differentiation.

**Figure 3.**
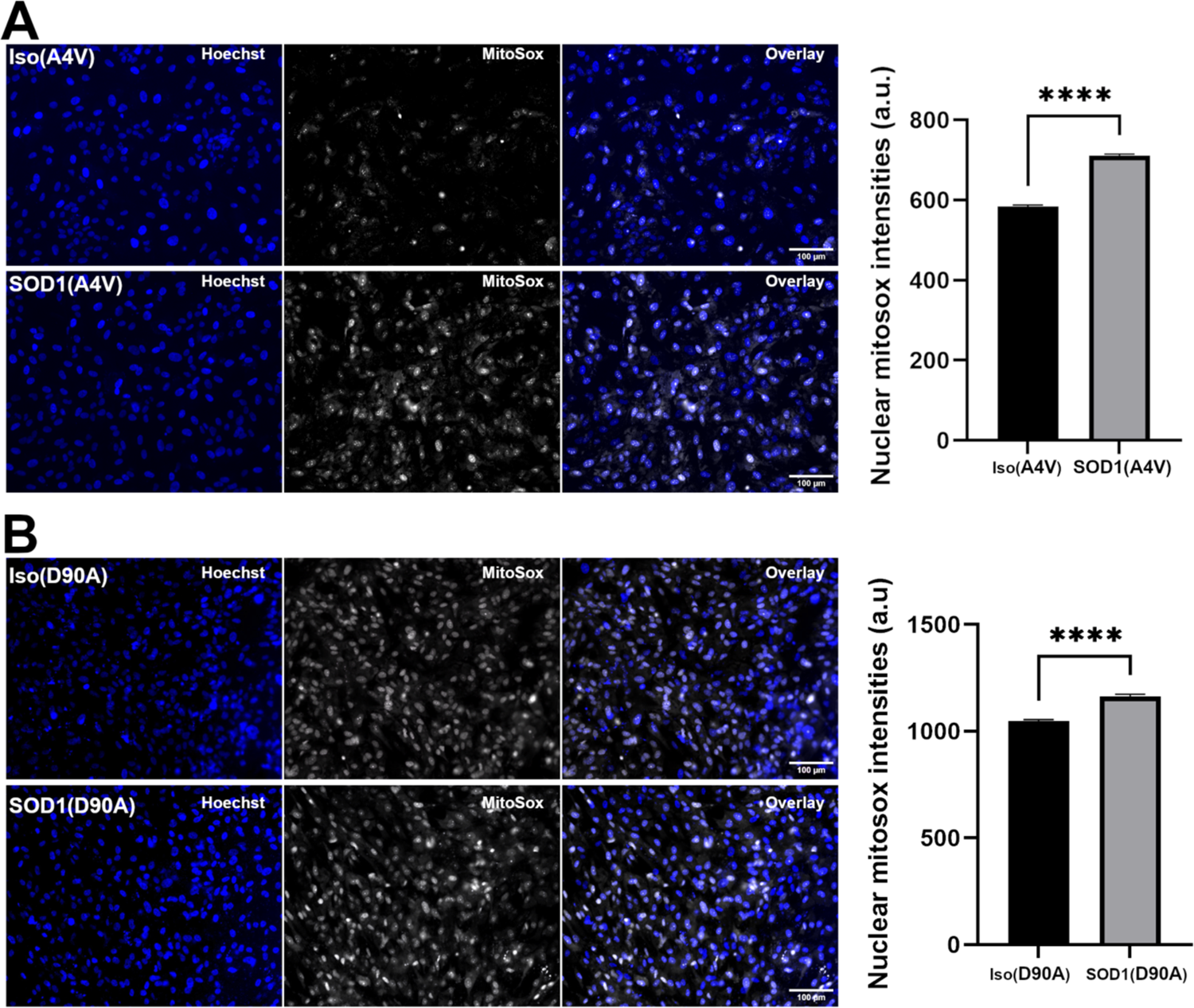
Nuclear oxidative stress in *SOD1*-mutated astrocytes. **A, B**) Representative images of nuclear MitoSox (5 μM) fluorescence in either *SOD1*(A4V) and isogenic astrocytes (A) or *SOD1*(D90A) and isogenic astrocytes (B). Graphs depict quantifications of nuclear MitoSox intensities in *SOD1*- mutated astrocytes compared to matching isogenic astrocytes. Statistical analyses were performed with Student’s t-test, graphs show mean ± *SEM*; \*\*\**p* < 0.0005; \*\*\*\**p*< 0.0001; *n* = 3 (more than 5,000 cells counted).

### 2.4. Increased DNA damage and nuclear accumulation of SOD1 protein in *SOD1*-mutated astrocytes

Considering the established link between oxidative stress and DNA damage in ALS (Kok et al., 2021; Szebényi et al., 2021), we next sought to determine whether the increased nuclear oxidative stress observed in *SOD1*-mutated astrocytes was correlated with increased DNA damage. To this end, we compared the levels of γH2AX, which represents the phosphorylated form of the histone variant H2AX and is a marker for DNA double strand breaks, in *SOD1*(A4V) and *SOD1*(D90A) astrocytes, compared to their isogenic counterparts. These studies revealed no detectable differences at DIV30 (not shown), but by DIV60 we observed an increased γH2AX signal in the nuclei of *SOD1*(A4V) and *SOD1*(D90A) astrocytes, indicative of increased double strand DNA break (Fig. 4A, B).

**Figure 4.**
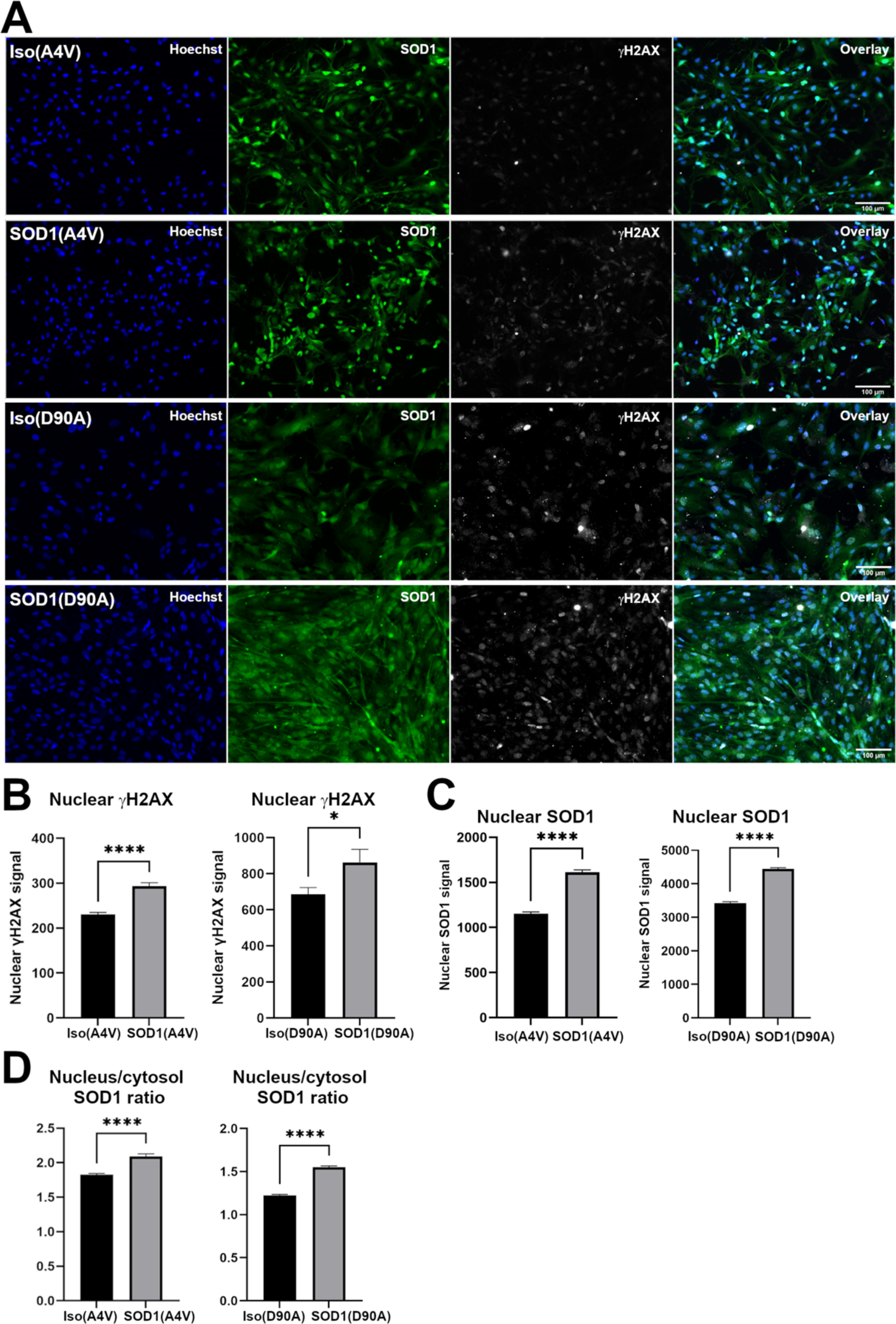
Increased DNA damage and SOD1 nuclear accumulation in SOD1-mutated astrocytes. **A)** Representative images of double-labeling immunofluorescence analysis of γH2AX and SOD1 in DIV60 *SOD1*(A4V) and *SOD1*(D90A) astrocytes together with their matching isogenic controls. **B-D)** Quantification of nuclear γH2AX (B), nuclear SOD1 (C), and nucleus vs cytosol SOD1 ratio (D) in DIV60 astrocytes of all four genotypes under study. Statistical analyses were performed with Student’s t-test, graphs show mean ± *SEM*; \**p*< 0.05; \*\*\**p* < 0.0005; \*\*\*\**p*< 0.0001; *n* = 3 (more than 5,000 cells counted).

Previous studies have shown that SOD1 accumulates in the nucleus in response to increased levels of reactive oxygen species and DNA damage (Inoue et al., 2010; Tsang et al., 2014). Moreover, an increase in nuclear versus cytosolic SOD1 protein localization was observed in ALS and other neurodegenerative disorders (Gertz et al., 2012; Bordoni et al., 2019). In the nucleus, in addition to its superoxide dismutase function, SOD1 acts a transcription factor that regulates the expression of oxidative resistance and DNA repair genes (Inoue et al., 2010; Tsang et al., 2014). Thus, we examined the intracellular localization of the SOD1 protein in mutated and isogenic human iPSC-derived astrocytes, both at earlier (DIV30) and later (DIV60) stages of *in vitro* differentiation. We observed no notable difference in nuclear SOD1 localization in *SOD1*(A4V) and *SOD1*(D90A) astrocytes compared to their isogenic counterparts at DIV30 (not shown). In contrast, a significant increase in nuclear SOD1, as detected through quantification of SOD1 signal within the nucleus, was observed in both *SOD1*(A4V) and *SOD1*(D90A) astrocytes at DIV60 (Fig. 4C). This observation was supported by quantification of the nucleus/cytosol SOD1 ratio in *SOD1*-mutated and isogenic astrocytes (Fig. 4D).

Taken together, these results suggest that astrocytes carrying ALS-associated *SOD1* mutations undergo early nuclear oxidative stress, which is then correlated with increased DNA damage, nuclear accumulation of SOD1, and reactive transformation at later *in vitro* stages.

## 3. Discussion

In this study, we sought to investigate the involvement of astrocytes in ALS, with specific focus on astrocytes harbouring *SOD1* mutations. Although rodent models have provided important insight into the contribution of astrocytes to ALS pathophysiology, there is growing evidence that human and murine astrocytes differ at various levels, including morphology, function, and expression of genes enriched in disease-associated pathways (Zhang et al., 2016; Kelley et al., 2018; Hodge et al., 2019). Moreover, most animal models are based on overexpression paradigms that do not fully recapitulate the pathophysiological expression levels occurring in human patients. The use of astrocytes generated from iPSCs derived from ALS patients carrying mutations in the *SOD1* gene can address both of these limitations, while also allowing the study of the impact of *SOD1* mutations on astrocyte biology in the absence of other cell type like MNs and microglia.

Using human astrocytes generated from iPSCs derived from familial ALS patients with *SOD1*(A4V) and *SOD1*(D90A) mutations we observed that these cells acquire a reactive phenotype during the first 60 days of *in vitro* culture. This cell-autonomous transformation agrees with previous studies showing increased reactivation of astrocytes harbouring mutations in several familial ALS genes (Birger et al., 2019; Taha et al., 2022; Stoklund Dittlau et al., 2023). Under the experimental conditions used in our investigations, *SOD1*-mutated astrocytes exhibited a “mixed” gene expression profile characterized by upregulation of both neurotoxic (A1 subtype) markers, such as *C1s*, *C3*, *SERPING1*, CXCL10, and neuroprotective (A2 subtype) markers, such as *PTX3, S100A10,* CCL2 (Liddelow et al., 2017). This finding agrees with previous studies showing up-regulation of both A1 and A2 marker genes in *SOD1*(D90A)-mutated astrocytes (Taha et al., 2022). These observations suggest that, in the absence of MNs and other cells implicated in ALS pathophysiology, such as microglia, astrocytes with mutations in *SOD1* are reactive but not neuroinflammatory under *in vitro* culture conditions.

In an effort to improve our understanding of the molecular mechanisms underlying the cell-autonomous reactive transformation of *SOD1*-mutated astrocytes, we observed that oxidative stress is detectable in the nuclei of these cells before overt signs of astrogliosis. To our knowledge, this is the first observation of increased levels of reactive oxygen species in the nucleus of astrocytes harbouring ALS mutations, and one of the earliest cell-autonomous phenotypes detected in these cells thus far. This nuclear phenotype was correlated with increased DNA damage, as well as nuclear accumulation of the SOD1 protein, at later *in vitro* stages.

These findings are consistent with previous evidence in multiple cell types that SOD1 becomes increasingly localized in the nucleus in response to both oxidative stress and DNA damage (Inoue et al., 2010; Tsang et al., 2014). The observation of nuclear oxidative stress before detection of DNA damage and SOD1 accumulation in the nucleus suggests a temporal sequence of nuclear phenotypes triggered by early nuclear oxidative damage. It remains to be determined whether it is the latter, or the ensuing DNA damage (or both), that contributes to SOD1 nuclear accumulation. SOD1 was previously detected in the nuclei of ventral horn astrocytes in *post-mortem* samples from ALS patients carrying *SOD1* mutations (Forsberg et al., 2011). Those studies could not determine whether this phenotype was dependent on the presence of MNs or other glial cells. Based on our findings, we propose that SOD1 nuclear accumulation in astrocytes harbouring ALS-associated *SOD1* mutations is a previously-unrecognized cell-autonomous mechanism. It is reasonable to assume that the increased presence of wild-type SOD1 in the nucleus would normally provide an early antioxidant and/or DNA repair function beneficial to astrocytes. The presence of mutated SOD1, however, would interfere with the physiological function of wild-type SOD1, leaving the cells more vulnerable to oxidative stress and DNA damage.

The nuclear phenotypes discussed above could be contributing factors to the increased reactivity of *SOD1*-mutated astrocytes compared to their isogenic counterparts. Oxidative stress in ALS *SOD1*-mutated astrocytes is involved in the neurotoxic effects of these cells on MN, as suggested by the observation that enhanced resistance to oxidative stress through increased levels of either total NAD content or SIRT6 protein can abrogate astrocyte toxicity toward co-cultured MNs (Harlan et al., 2019). This finding is consistent with results showing that astrocytes derived from iPSCs from ALS patients with mutated *C9orf72* exhibit increased oxidative stress and neurotoxicity (Birger et al., 2019). Moreover, analysis of *post-mortem* tissue from sporadic ALS patients revealed that poly(ADP-ribose) polymerase, a key DNA repair protein, is increased in astrocytes, suggesting increased DNA damage in astrocytes in ALS (Kim et al., 2003). Consistently, DNA damage response pathways are affected in both astroglia and neurons in brain organoid slice cultures derived from iPSCs with mutated *C9orf72* (Szebényi et al., 2021).

Additionally, evidence that DNA damage can contribute to astrocyte dysfunction in neurodegenerative diseases has recently emerged from the study of astrocytes from Huntington’s Disease patients (Lange et al., 2023). Perturbation of nuclear SOD1 functions is also expected to contribute to enhanced astrocyte reactivation. SOD1 localizes to the nucleus under normal and pathological conditions to contribute to oxidative stress response and DNA repair mechanisms (Inoue et al., 2010; Tsang et al., 2014; Xu et al., 2022). It is therefore reasonable to hypothesize that a dominant-negative effect of mutated SOD1 on the nuclear roles of wild-type SOD1 would lead to increased oxidative stress and DNA damage in astrocytes, thereby contributing to mechanisms promoting astrogliosis.

In summary, the present study has characterized early cell-autonomous mechanisms of astrocyte dysfunction associated with two of the most prevalent *SOD1* mutations in ALS patients. The described astrocyte phenotypes have the potential to contribute to mechanisms of MN degeneration in ALS, further underscoring the importance of considering astrocyte dysfunction when developing therapies for ALS.

## 4. Materials and Methods

### 4.1. Human induced pluripotent stem cells

Human iPSC lines *SOD1*(A4V) and *SOD1*(D90A) were obtained from Target ALS (https://www.targetals.org; Cat No. ND35671 (A4V) and ND35660 (D90A). To generate matching isogenic control iPSC lines Iso(A4V) and Iso(D90A) from the parental lines, CRISPR editing was performed using established methods (Deneault et al., 2021). All iPSC lines were maintained at the Montreal Neurological Institute-Hospital through procedures conducted under Ethical Review Board approval by the McGill University Health Centre Board (DURCAN_IPSC/2019-5374). Undifferentiated state of iPSCs was assessed by testing for expression of the stem cell markers NANOG and OCT4 using rabbit anti-NANOG (1/1,000; Abcam; Cambridge, UK; Cat. No. ab21624) and rabbit anti-OCT4 (1 μg/ml; Abcam; Cat. No. ab19857) or goat anti-OCT3/4 (1/500; Santa Cruz Biotechnology; Dallas, TX, USA, Cat. No. sc-8628) antibodies, and by quality control profiling as described previously (Chen et al., 2021).

### 4.2. Derivation of neural progenitor cells from human iPSCs

Human iPSCs at low passage number were cultured in mTeSR medium (STEMCELL Technologies; Vancouver, BC, Canada; Cat. No. 85850) in 10-cm culture dishes (Thermo-Fisher Scientific; Waltham, MA; Cat. No. 353003) coated with Matrigel (Thermo-Fisher Scientific; Cat. No. 08-774-552) until they reached 70%-80% confluence. To generate NPCs, iPSCs were dissociated with Gentle Cell Dissociation Reagent (STEMCELL Technologies; Cat. No. 07174), followed by seeding of 2-3×10^6^ cells onto T25 flasks (Thermo-Fisher Scientific; Cat. No. 12-556-009) coated with Matrigel. Cells were then cultured overnight with 5 ml mTeSR supplemented with 10 μM ROCK inhibitor (compound Y-27632 2HCl; Selleck Chemicals; Houston, TX, USA; Cat. No. S1049). At *in vitro* day 1 (DIV1), mTeSR was replaced with ‘neural induction medium’ containing DMEM/F12 supplemented with GlutaMax (1/1; Thermo-Fisher Scientific; Cat. No. 10565-018), Neurobasal medium (1/1; Thermo-Fisher Scientific; Cat. No. 21103-049), N2 (0.5X; Thermo-Fisher Scientific; Cat. No. 17504-044), B27 (0.5X; Thermo-Fisher Scientific; Cat. No. 17502-048), ascorbic acid (100 μM; Sigma-Aldrich; St. Louis, MO, USA; Cat. No. A5960), L-Glutamax (0.5X; Thermo-Fisher Scientific; Cat. No. 35050-061), antibiotic-antimycotic (1X; Thermo-Fisher Scientific; Cat. No. 15240-062), 3 μM CHIR99021 (STEMCELL Technologies; Cat. No. 72054), 2 μM DMH1 (Sigma-Aldrich; Cat. No. D8946), and 2 μM SB431542 (Tocris Bioscience; Bristol, UK; Cat. No. 1614). The culture medium was changed every other day until DIV6, when induced NPCs were instructed to acquire a caudalized and ventralized progenitor cell identity as follows. NPCs were dissociated with Gentle Cell Dissociation Reagent and split 1:6 with the same medium described above, supplemented with retinoic acid (RA) (0.1 μM; Sigma-Aldrich; Cat. No. R2625) and purmorphamine (0.5 μM; Sigma-Aldrich; Cat. No. SML-0868) in combination with 1 μM CHIR99021, 2 μM DMH1 and 2 μM SB431542 reagents. The culture medium was changed every other day until DIV12, when cells were split again 1:6 and expanded with the same medium containing 3 μM CHIR99021, 2 μM DMH1, 2 μM SB431542, 0.1 μM RA, 0.5 μM purmorphamine, and 500 μM valproic acid (VPA; Sigma-Aldrich; Cat. No. P4543) till DIV18. The ensuing caudalized and ventralized NPCs were validated by real-time polymerase chain reaction (RT-PCR) and immunocytochemistry.

### 4.3. Differentiation of astrocytes from human iPSC-derived neural progenitor cells

Induced caudalized/ventralized NPCs were differentiated into astrocytes starting at DIV18 using a defined medium. NPCs were seeded at low cell density (15,000 cells/cm^2^) in two T25 flasks in the presence of 5 ml of NPC expansion medium containing ROCK inhibitor. Next day, medium was replaced with ‘Astrocyte Differentiation Medium 1’ [ScienceCell Astrocyte Growth Medium (ScienCell Research Laboratories; Carlsbad, CA, USA; Cat. No. 1801b) containing astrocyte growth supplement (ScienCell Research Laboratories; Cat. No. 1852), 1% fetal bovine serum (FBS) (ScienCell Research Laboratories; Cat. No. 0010), 50 U/ml penicillin G, 50 mg/ml streptomycin]. Cells were split 1:4 every week and maintained under these culture conditions for 30 days. Half medium was replaced with fresh medium every 3 to 4 days. At DIV50, cultures were switched to ‘Astrocyte Differentiation Medium 2’ (same as Astrocyte Differentiation Medium 1 but lacking FBS). Induced astrocytes were validated by immunocytochemistry, RT-PCR, and by measuring their response to treatment with a cocktail of IL-1α, TNF-α, and C1q (Liddelow et al., 2017).

### 4.4. Characterization of induced cells by immunocytochemistry

Induced human NPCs and astrocytes were analyzed by immunocytochemistry, which was performed as described previously (Methot et al., 2018). The following primary antibodies were used: rabbit anti HOXA5 (1/67000; kindly provided by Dr. Jeremy Dasen, New York, University School of Medicine), mouse anti-NKX6.1 (1/500; DSHB; Iowa City, IA; Cat. No. F55A10), mouse anti-GFAP (1/1,000; Sigma-Aldrich; Cat. No. G3893); mouse anti-S100B (1/500; Sigma-Aldrich; Cat. No. S2532), mouse anti-γH2AX (1/500; Millipore; Burlington, MA; Cat. No. 05636), rabbit anti-SOD1 (1/500; Enzo Life Sciences; Farmingdale, NY; Cat. No. ADI-SOD-100-F). Secondary antibodies against primary reagents raised in various species were conjugated to Alexa Fluor 555, Alexa Fluor 488 (1/1,000; Invitrogen; Burlington, ON, Canada). Actin polymerization was visualized by staining of F-actin using Alexa-Fluor-488 phalloidin (1/500; Thermo Fisher; Cat. No. A12379). Images were acquired with a Zeiss Axio Observer Z1 Inverted Microscope using 20X magnification (N.A 0.8) and a ZEISS Axiocam 506 mono camera.

### 4.5. Characterization of induced cells by real-time polymerase chain reaction

RNA extraction and real-time polymerase chain reaction (RT-PCR) were performed as described (Soubannier et al., 2020). Analysis of gene expression was conducted using the following oligonucleotide primers: Taqman probes *C1*s, Hs00156159_m1; *C3*, Hs00163811_m1; *CCL2*, Hs00234140_m1; *CXCL10*, Hs00171042_m1; *S100A10*, Hs00237010_m1; *PTX3*, Hs00173615_m1; *SERPING1*, Hs00163781_m1. Primer/probe sets were obtained from ThermoFisher Scientific. Data were normalized with *BETA-ACTIN* and *GAPDH* (*ACTB Hs01060665_g1; GAPDH Hs02786624_g1).* Relative quantification (RQ) was estimated according to the ΔCt method (Schmittgen and Livak, 2008).

### 4.6. Quantification of protein levels

Conditioned media collected from astrocytes treated, or not, with TNF-α (30 ng/ml), IL-1α (3 ng/ml), and C1q (400 ng/ml) for 48 hr were centrifuged at 300 x g for 5 min and supernatant was recovered. Supernatants were sent for proteomics analysis to SomaLogic Inc., Boulder, Colorado (https://somalogic.com/). For studies comparing conditioned media from cultures of Iso(A4V) and SOD1(A4V) astrocytes, protein levels were analyzed using Bio-Plex Pro Human Cytokine Screening Panel (48-Plex) (Cat. No. 12007283), Bio-Plex Pro Human Chemokine Panel (40-Plex) (Cat. No. 171AK99MR2), and Bio-Plex Pro Human Inflammation Panel 1 (37-plex) (Cat. No. 171AL001M) from Bio-Rad (Hercules, CA).

### 4.7. Quantification of reactive oxygen species

Cells were incubated in presence of either 1 μM or 5 μM MitoSox^TM^ (Invitrogen; Cat. #M36008) for 30 minutes in medium containing Hoechst (1/2,000 dilution). Following the incubation period, cells were washed for 5 minutes with 37°C-preheated astrocyte growth medium containing astrocyte growth supplement and 1% FBS. Subsequently, the cell culture medium was replaced once again, and the cells were subjected to microscopic observation. For nuclear MitoSox intensities quantification, regions of interest (ROIs) were obtained from the channel corresponding to the Hoechst staining through thresholding. Each ROI corresponding to the nucleus was then used to measure the mean intensity signal in the MitoSox fluorescence channel. Several pictures corresponding to at least 5,000 cells were counted.

## Authorship contribution statement

VS performed all cell culture and microscopy experiments, data analysis, and figure preparation. MC, LG, SL, DK, GH performed experiments. SS, TMD, VS, GR conceived overall study plan. SS, TMD supervised the study. SS and VS wrote the manuscript.

## Acknowledgments

We thank Anna Kristyna Franco Flores for experimental assistance, and Valerio Piscopo for discussions and advice.

## References

Abati E, Bresolin N, Comi G, Corti S. Silence superoxide dismutase 1 (SOD1): a promising therapeutic target for amyotrophic lateral sclerosis (ALS). Expert Opin Ther Targets. 2020 Apr;24(4):295–310. doi: 10.1080/14728222.2020.1738390. Epub 2020 Mar 14.

Appel SH, Beers DR, Zhao W. Amyotrophic lateral sclerosis is a systemic disease: peripheral contributions to inflammation-mediated neurodegeneration. Curr Opin Neurol. 2021 Oct 1;34(5):765–772. doi: 10.1097/WCO.0000000000000983.

Arnold FJ, Nguyen AD, Bedlack RS, Bennett CL, La Spada AR. Intercellular transmission of pathogenic proteins in ALS: Exploring the pathogenic wave. Neurobiol Dis. 2023 Jun 30;184:106218. doi: 10.1016/j.nbd.2023.106218. Epub ahead of print.

Arredondo C, Cefaliello C, Dyrda A, Jury N, Martinez P, Díaz I, Amaro A, Tran H, Morales D, Pertusa M, Stoica L, Fritz E, Corvalán D, Abarzúa S, Méndez-Ruette M, Fernández P, Rojas F, Kumar MS, Aguilar R, Almeida S, Weiss A, Bustos FJ, González-Nilo F, Otero C, Tevy MF, Bosco DA, Sáez JC, Kähne T, Gao FB, Berry JD, Nicholson K, Sena-Esteves M, Madrid R, Varela D, Montecino M, Brown RH, van Zundert B. Excessive release of inorganic polyphosphate by ALS/FTD astrocytes causes non-cell-autonomous toxicity to motoneurons. Neuron. 2022 May 18;110(10):1656–1670.e12. doi: 10.1016/j.neuron.2022.02.010. Epub 2022 Mar 10.

Balendra R, Isaacs AM. C9orf72-mediated ALS and FTD: multiple pathways to disease. Nat Rev Neurol. 2018 Sep;14(9):544–558. doi: 10.1038/s41582-018-0047-2.

Beers DR, Appel SH. Immune dysregulation in amyotrophic lateral sclerosis: mechanisms and emerging therapies. Lancet Neurol. 2019 Feb;18(2):211–220. doi: 10.1016/S1474-4422(18)30394-6.

Birger A, Ben-Dor I, Ottolenghi M, Turetsky T, Gil Y, Sweetat S, Perez L, Belzer V, Casden N, Steiner D, Izrael M, Galun E, Feldman E, Behar O, Reubinoff B. Human iPSC-derived astrocytes from ALS patients with mutated C9ORF72 show increased oxidative stress and neurotoxicity. EBioMedicine. 2019 Dec;50:274–289. doi: 10.1016/j.ebiom.2019.11.026. Epub 2019 Nov 29.

Bordoni M, Pansarasa O, Dell’Orco M, Crippa V, Gagliardi S, Sproviero D, Bernuzzi S, Diamanti L, Ceroni M, Tedeschi G, Poletti A, Cereda C. Nuclear Phospho-SOD1 Protects DNA from Oxidative Stress Damage in Amyotrophic Lateral Sclerosis. J Clin Med. 2019 May 22;8(5):729. doi: 10.3390/jcm8050729.

Brenner D, Freischmidt A. Update on genetics of amyotrophic lateral sclerosis. Curr Opin Neurol. 2022 Oct 1;35(5):672–677. doi: 10.1097/WCO.0000000000001093. Epub 2022 Aug 8.

Bunton-Stasyshyn RK, Saccon RA, Fratta P, Fisher EM. SOD1 Function and Its Implications for Amyotrophic Lateral Sclerosis Pathology: New and Renascent Themes. Neuroscientist. 2015 Oct;21(5):519–29. doi: 10.1177/1073858414561795. Epub 2014 Dec 9.

Burk K, Pasterkamp RJ. Disrupted neuronal trafficking in amyotrophic lateral sclerosis. Acta Neuropathol. 2019 Jun;137(6):859–877. doi: 10.1007/s00401-019-01964-7. Epub 2019 Feb 5.

Calabrese G, Molzahn C, Mayor T. Protein interaction networks in neurodegenerative diseases: From physiological function to aggregation. J Biol Chem. 2022 Jul;298(7):102062. doi: 10.1016/j.jbc.2022.102062. Epub 2022 May 25.

Chen CX, Abdian N, Maussion G, Thomas RA, Demirova I, Cai E, Tabatabaei M, Beitel LK, Karamchandani J, Fon EA, Durcan TM. A Multistep Workflow to Evaluate Newly Generated iPSCs and Their Ability to Generate Different Cell Types. Methods Protoc. 2021 Jul 19;4(3):50. doi: 10.3390/mps4030050.

Cipollina G, Davari Serej A, Di Nolfi G, Gazzano A, Marsala A, Spatafora MG, Peviani M. Heterogeneity of Neuroinflammatory Responses in Amyotrophic Lateral Sclerosis: A Challenge or an Opportunity? Int J Mol Sci. 2020 Oct 25;21(21):7923. doi: 10.3390/ijms21217923.

Deneault E, Chaineau M, Nicouleau M, Castellanos Montiel MJ, Franco Flores AK, Haghi G, Chen CX, Abdian N, Shlaifer I, Beitel LK, Durcan TM. A streamlined CRISPR workflow to introduce mutations and generate isogenic iPSCs for modeling amyotrophic lateral sclerosis. Methods. 2022 Jul;203:297–310. doi: 10.1016/j.ymeth.2021.09.002. Epub 2021 Sep 6.

Di Giorgio FP, Carrasco MA, Siao MC, Maniatis T, Eggan K. Non-cell autonomous effect of glia on motor neurons in an embryonic stem cell-based ALS model. Nat Neurosci. 2007 May;10(5):608–14. doi: 10.1038/nn1885. Epub 2007 Apr 15.

Forsberg K, Andersen PM, Marklund SL, Brännström T. Glial nuclear aggregates of superoxide dismutase-1 are regularly present in patients with amyotrophic lateral sclerosis. Acta Neuropathol. 2011 May;121(5):623–34. doi: 10.1007/s00401-011-0805-3. Epub 2011 Feb 3.

Fritz E, Izaurieta P, Weiss A, Mir FR, Rojas P, Gonzalez D, Rojas F, Brown RH Jr, Madrid R, van Zundert B. Mutant SOD1-expressing astrocytes release toxic factors that trigger motoneuron death by inducing hyperexcitability. J Neurophysiol. 2013 Jun;109(11):2803–14. doi: 10.1152/jn.00500.2012. Epub 2013 Mar 13.

Gertz B, Wong M, Martin LJ. Nuclear localization of human SOD1 and mutant SOD1-specific disruption of survival motor neuron protein complex in transgenic amyotrophic lateral sclerosis mice. J Neuropathol Exp Neurol. 2012 Feb;71(2):162–77. doi: 10.1097/NEN.0b013e318244b635.

Guttenplan KA, Weigel MK, Adler DI, Couthouis J, Liddelow SA, Gitler AD, Barres BA. Knockout of reactive astrocyte activating factors slows disease progression in an ALS mouse model. Nat Commun. 2020 Jul 27;11(1):3753. doi: 10.1038/s41467-020-17514-9.

Hall ED, Oostveen JA, Gurney ME. Relationship of microglial and astrocytic activation to disease onset and progression in a transgenic model of familial ALS. Glia. 1998 Jul;23(3):249–56. doi: 10.1002/(sici)1098-1136(199807)23:3<249::aid-glia7>3.0.co;2-#.

Haidet-Phillips AM, Hester ME, Miranda CJ, Meyer K, Braun L, Frakes A, Song S, Likhite S, Murtha MJ, Foust KD, Rao M, Eagle A, Kammesheidt A, Christensen A, Mendell JR, Burghes AH, Kaspar BK. Astrocytes from familial and sporadic ALS patients are toxic to motor neurons. Nat Biotechnol. 2011 Aug 10;29(9):824–8. doi: 10.1038/nbt.1957.

Hansson E. Actin filament reorganization in astrocyte networks is a key functional step in neuroinflammation resulting in persistent pain: novel findings on network restoration. Neurochem Res. 2015 Feb;40(2):372–9. doi: 10.1007/s11064-014-1363-6. Epub 2014 Jun 21.

Harlan BA, Pehar M, Killoy KM, Vargas MR. Enhanced SIRT6 activity abrogates the neurotoxic phenotype of astrocytes expressing ALS-linked mutant SOD1. FASEB J. 2019 Jun;33(6):7084–7091. doi: 10.1096/fj.201802752R. Epub 2019 Mar 6.

Hodge RD, Bakken TE, Miller JA, Smith KA, Barkan ER, Graybuck LT, Close JL, Long B, Johansen N, Penn O, Yao Z, Eggermont J, Höllt T, Levi BP, Shehata SI, Aevermann B, Beller A, Bertagnolli D, Brouner K, Casper T, Cobbs C, Dalley R, Dee N, Ding SL, Ellenbogen RG, Fong O, Garren E, Goldy J, Gwinn RP, Hirschstein D, Keene CD, Keshk M, Ko AL, Lathia K, Mahfouz A, Maltzer Z, McGraw M, Nguyen TN, Nyhus J, Ojemann JG, Oldre A, Parry S, Reynolds S, Rimorin C, Shapovalova NV, Somasundaram S, Szafer A, Thomsen ER, Tieu M, Quon G, Scheuermann RH, Yuste R, Sunkin SM, Lelieveldt B, Feng D, Ng L, Bernard A, Hawrylycz M, Phillips JW, Tasic B, Zeng H, Jones AR, Koch C, Lein ES. Conserved cell types with divergent features in human versus mouse cortex. Nature. 2019 Sep;573(7772):61-68. doi: 10.1038/s41586-019-1506-7. Epub 2019 Aug 21.

Inoue E, Tano K, Yoshii H, Nakamura J, Tada S, Watanabe M, Seki M, Enomoto T. SOD1 Is Essential for the Viability of DT40 Cells and Nuclear SOD1 Functions as a Guardian of Genomic DNA. J Nucleic Acids. 2010 Aug 5;2010:795946. doi: 10.4061/2010/795946.

Johann S, Heitzer M, Kanagaratnam M, Goswami A, Rizo T, Weis J, Troost D, Beyer C. NLRP3 inflammasome is expressed by astrocytes in the SOD1 mouse model of ALS and in human sporadic ALS patients. Glia. 2015 Dec;63(12):2260–73. doi: 10.1002/glia.22891.

Kelley KW, Nakao-Inoue H, Molofsky AV, Oldham MC. Variation among intact tissue samples reveals the core transcriptional features of human CNS cell classes. Nat Neurosci. 2018 Sep;21(9):1171–1184. doi: 10.1038/s41593-018-0216-z. Epub 2018 Aug 28.

Kim SH, Henkel JS, Beers DR, Sengun IS, Simpson EP, Goodman JC, Engelhardt JI, Siklós L, Appel SH. PARP expression is increased in astrocytes but decreased in motor neurons in the spinal cord of sporadic ALS patients. J Neuropathol Exp Neurol. 2003 Jan;62(1):88–103. doi: 10.1093/jnen/62.1.88.

Kim G, Gautier O, Tassoni-Tsuchida E, Ma XR, Gitler AD. ALS Genetics: Gains, Losses, and Implications for Future Therapies. Neuron. 2020 Dec 9;108(5):822–842. doi: 10.1016/j.neuron.2020.08.022. Epub 2020 Sep 14.

Kok JR, Palminha NM, Dos Santos Souza C, El-Khamisy SF, Ferraiuolo L. DNA damage as a mechanism of neurodegeneration in ALS and a contributor to astrocyte toxicity. Cell Mol Life Sci. 2021 Aug;78(15):5707–5729. doi: 10.1007/s00018-021-03872-0.

Lange J, Gillham O, Flower M, Ging H, Eaton S, Kapadia S, Neueder A, Duchen MR, Ferretti P, Tabrizi SJ. PolyQ length-dependent metabolic alterations and DNA damage drive human astrocyte dysfunction in Huntington’s disease. Prog Neurobiol. 2023 Jun;225:102448. doi: 10.1016/j.pneurobio.2023.102448. Epub 2023 Apr 5.

Liddelow SA, Guttenplan KA, Clarke LE, Bennett FC, Bohlen CJ, Schirmer L, Bennett ML, Münch AE, Chung WS, Peterson TC, Wilton DK, Frouin A, Napier BA, Panicker N, Kumar M, Buckwalter MS, Rowitch DH, Dawson VL, Dawson TM, Stevens B, Barres BA. Neurotoxic reactive astrocytes are induced by activated microglia. Nature. 2017 Jan 26;541(7638):481-487. doi: 10.1038/nature21029. Epub 2017 Jan 18.

Marchetto MC, Muotri AR, Mu Y, Smith AM, Cezar GG, Gage FH. Non-cell-autonomous effect of human SOD1 G37R astrocytes on motor neurons derived from human embryonic stem cells. Cell Stem Cell. 2008 Dec 4;3(6):649–57. doi: 10.1016/j.stem.2008.10.001.

Mejzini R, Flynn LL, Pitout IL, Fletcher S, Wilton SD, Akkari PA. ALS Genetics, Mechanisms, and Therapeutics: Where Are We Now? Front Neurosci. 2019 Dec 6;13:1310. doi: 10.3389/fnins.2019.01310.

Methot L, Soubannier V, Hermann R, Campos E, Li S, Stifani S. Nuclear factor-kappaB regulates multiple steps of gliogenesis in the developing murine cerebral cortex. Glia. 2018 Dec;66(12):2659–2672. doi: 10.1002/glia.23518. Epub 2018 Oct 19.

Meyer K, Ferraiuolo L, Miranda CJ, Likhite S, McElroy S, Renusch S, Ditsworth D, Lagier-Tourenne C, Smith RA, Ravits J, Burghes AH, Shaw PJ, Cleveland DW, Kolb SJ, Kaspar BK. Direct conversion of patient fibroblasts demonstrates non-cell autonomous toxicity of astrocytes to motor neurons in familial and sporadic ALS. Proc Natl Acad Sci U S A. 2014 Jan 14;111(2):829–32. doi: 10.1073/pnas.1314085111. Epub 2013 Dec 30.

Nagai M, Re DB, Nagata T, Chalazonitis A, Jessell TM, Wichterle H, Przedborski S. Astrocytes expressing ALS-linked mutated SOD1 release factors selectively toxic to motor neurons. Nat Neurosci. 2007 May;10(5):615–22. doi: 10.1038/nn1876. Epub 2007 Apr 15.

Okado-Matsumoto A, Fridovich I. Subcellular distribution of superoxide dismutases (SOD) in rat liver: Cu,Zn-SOD in mitochondria. J Biol Chem. 2001 Oct 19;276(42):38388–93. doi: 10.1074/jbc.M105395200. Epub 2001 Aug 15.

Peggion C, Scalcon V, Massimino ML, Nies K, Lopreiato R, Rigobello MP, Bertoli A. SOD1 in ALS: Taking Stock in Pathogenic Mechanisms and the Role of Glial and Muscle Cells. Antioxidants (Basel). 2022 Mar 23;11(4):614. doi: 10.3390/antiox11040614.

Prasad A, Bharathi V, Sivalingam V, Girdhar A, Patel BK. Molecular Mechanisms of TDP-43 Misfolding and Pathology in Amyotrophic Lateral Sclerosis. Front Mol Neurosci. 2019 Feb 14;12:25. doi: 10.3389/fnmol.2019.00025.

Roelofs BA, Ge SX, Studlack PE, Polster BM. Low micromolar concentrations of the superoxide probe MitoSOX uncouple neural mitochondria and inhibit complex IV. Free Radic Biol Med. 2015 Sep;86:250–8. doi: 10.1016/j.freeradbiomed.2015.05.032. Epub 2015 Jun 6.

Rosen DR, Siddique T, Patterson D, Figlewicz DA, Sapp P, Hentati A, Donaldson D, Goto J, O’Regan JP, Deng HX, et al. Mutations in Cu/Zn superoxide dismutase gene are associated with familial amyotrophic lateral sclerosis. Nature. 1993 Mar 4;362(6415):59-62. doi: 10.1038/362059a0.

Schiffer D, Cordera S, Cavalla P, Migheli A. Reactive astrogliosis of the spinal cord in amyotrophic lateral sclerosis. J Neurol Sci. 1996 Aug;139 Suppl:27-33. doi: 10.1016/0022-510x(96)00073-1.

Schmittgen TD, Livak KJ. Analyzing real-time PCR data by the comparative C(T) method. Nat Protoc. 2008;3(6):1101–8. doi: 10.1038/nprot.2008.73

Shibata N, Nagai R, Uchida K, Horiuchi S, Yamada S, Hirano A, Kawaguchi M, Yamamoto T, Sasaki S, Kobayashi M. Morphological evidence for lipid peroxidation and protein glycoxidation in spinal cords from sporadic amyotrophic lateral sclerosis patients. Brain Res. 2001 Oct 26;917(1):97–104. doi: 10.1016/s0006-8993(01)02926-2.

Stoklund Dittlau K, Terrie L, Baatsen P, Kerstens A, De Swert L, Janky R, Corthout N, Masrori P, Van Damme P, Hyttel P, Meyer M, Thorrez L, Freude K, Van Den Bosch L. FUS-ALS hiPSC-derived astrocytes impair human motor units through both gain-of-toxicity and loss-of-support mechanisms. Mol Neurodegener. 2023 Jan 18;18(1):5. doi: 10.1186/s13024-022-00591-3.

Soubannier V, Maussion G, Chaineau M, Sigutova V, Rouleau G, Durcan TM, Stifani S. Characterization of human iPSC-derived astrocytes with potential for disease modeling and drug discovery. Neurosci Lett. 2020 Jul 13;731:135028. doi: 10.1016/j.neulet.2020.135028. Epub 2020 May 4.

Soubannier V, Chaineau M, Gursu L, Haghi G, Franco Flores AK, Rouleau G, Durcan TM, Stifani S. Rapid Generation of Ventral Spinal Cord-like Astrocytes from Human iPSCs for Modeling Non-Cell Autonomous Mechanisms of Lower Motor Neuron Disease. Cells. 2022 Jan 24;11(3):399. doi: 10.3390/cells11030399.

Szebényi K, Wenger LMD, Sun Y, Dunn AWE, Limegrover CA, Gibbons GM, Conci E, Paulsen O, Mierau SB, Balmus G, Lakatos A. Human ALS/FTD brain organoid slice cultures display distinct early astrocyte and targetable neuronal pathology. Nat Neurosci. 2021 Nov;24(11):1542–1554. doi: 10.1038/s41593-021-00923-4. Epub 2021 Oct 21.

Taha DM, Clarke BE, Hall CE, Tyzack GE, Ziff OJ, Greensmith L, Kalmar B, Ahmed M, Alam A, Thelin EP, Garcia NM, Helmy A, Sibley CR, Patani R. Astrocytes display cell autonomous and diverse early reactive states in familial amyotrophic lateral sclerosis. Brain. 2022 Apr 18;145(2):481–489. doi: 10.1093/brain/awab328.

Tran NN, Lee BH. Functional implication of ubiquitinating and deubiquitinating mechanisms in TDP-43 proteinopathies. Front Cell Dev Biol. 2022 Sep 9;10:931968. doi: 10.3389/fcell.2022.931968.

Tsang CK, Liu Y, Thomas J, Zhang Y, Zheng XF. Superoxide dismutase 1 acts as a nuclear transcription factor to regulate oxidative stress resistance. Nat Commun. 2014 Mar 19;5:3446. doi: 10.1038/ncomms4446.

Tyzack GE, Hall CE, Sibley CR, Cymes T, Forostyak S, Carlino G, Meyer IF, Schiavo G, Zhang SC, Gibbons GM, Newcombe J, Patani R, Lakatos A. A neuroprotective astrocyte state is induced by neuronal signal EphB1 but fails in ALS models. Nat Commun. 2017 Oct 27;8(1):1164. doi: 10.1038/s41467-017-01283-z.

Urban MW, Charsar BA, Heinsinger NM, Markandaiah SS, Sprimont L, Zhou W, Henderson NT, Ghosh B, Cain RE, Trotti D, Pasinelli P, Wright MC, Dalva MB, Lepore AC. EphrinB2 knockdown in spinal cord astrocytes preserves diaphragm innervation in a mutant SOD1 mouse model of ALS. bioRxiv [Preprint]. 2023 May 10:2023.05.10.538887. doi: 10.1101/2023.05.10.538887.

Van Harten ACM, Phatnani H, Przedborski S. Non-cell-autonomous pathogenic mechanisms in amyotrophic lateral sclerosis. Trends Neurosci. 2021 Aug;44(8):658–668. doi: 10.1016/j.tins.2021.04.008. Epub 2021 May 15.

Varcianna A, Myszczynska MA, Castelli LM, O’Neill B, Kim Y, Talbot J, Nyberg S, Nyamali I, Heath PR, Stopford MJ, Hautbergue GM, Ferraiuolo L. Micro-RNAs secreted through astrocyte-derived extracellular vesicles cause neuronal network degeneration in C9orf72 ALS. EBioMedicine. 2019 Feb;40:626–635. doi: 10.1016/j.ebiom.2018.11.067. Epub 2019 Jan 31.

Xu J, Su X, Burley SK, Zheng XFS. Nuclear SOD1 in Growth Control, Oxidative Stress Response, Amyotrophic Lateral Sclerosis, and Cancer. Antioxidants (Basel). 2022 Feb 21;11(2):427. doi: 10.3390/antiox11020427.

Zhang Y, Sloan SA, Clarke LE, Caneda C, Plaza CA, Blumenthal PD, Vogel H, Steinberg GK, Edwards MS, Li G, Duncan JA 3rd, Cheshier SH, Shuer LM, Chang EF, Grant GA, Gephart MG, Barres BA. Purification and Characterization of Progenitor and Mature Human Astrocytes Reveals Transcriptional and Functional Differences with Mouse. Neuron. 2016 Jan 6;89(1):37–53. doi: 10.1016/j.neuron.2015.11.013. Epub 2015 Dec 10.

Zhou Y, Liu X, Ma S, Zhang N, Yang D, Wang L, Ye S, Zhang Q, Ruan J, Ma J, Wang S, Jiang N, Zhao Z, Zhao S, Zheng C, Fan X, Gong Y, Abdoul Razak MY, Hu W, Pan J, Wang X, Fan J, Li J, Liu R, Shentu Y. ChK1 activation induces reactive astrogliosis through CIP2A/PP2A/STAT3 pathway in Alzheimer’s disease. FASEB J. 2022 Mar;36(3):e22209. doi: 10.1096/fj.202101625R.

