## Supplementary Figures for "Early nuclear phenotypes and reactive transformation in human iPSC-derived astrocytes from ALS patients with *SOD1* mutations"

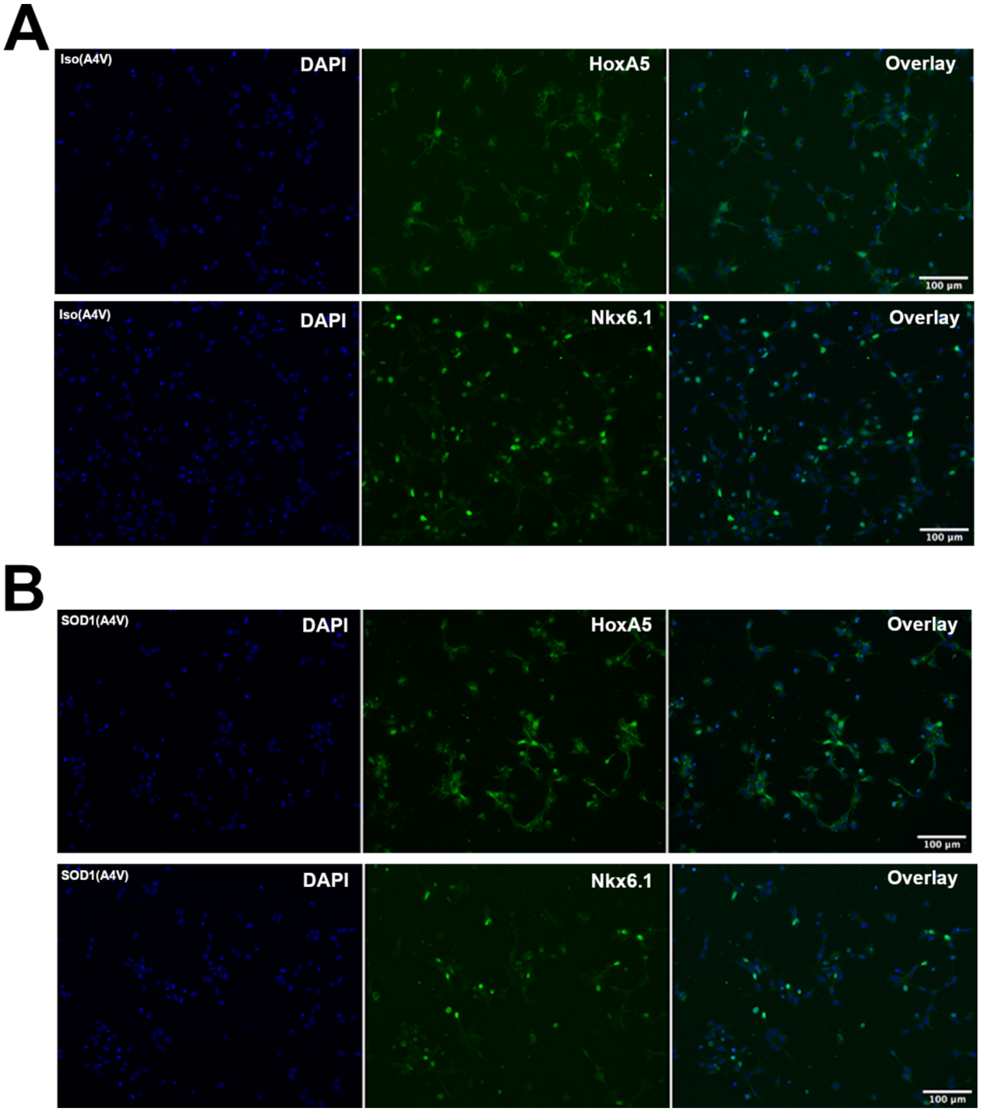


**Supplementary Figure 1. A, B**) Representative images of immunofluorescence analysis of HOXA5 and NKX6.1 (green) expression in neural progenitor cells generated from *SOD1*(A4V) mutation-corrected, Iso(A4V) (A) and *SOD1*(A4V) mutated (B) iPSC lines; Hoechst counterstaining (blue) is shown.

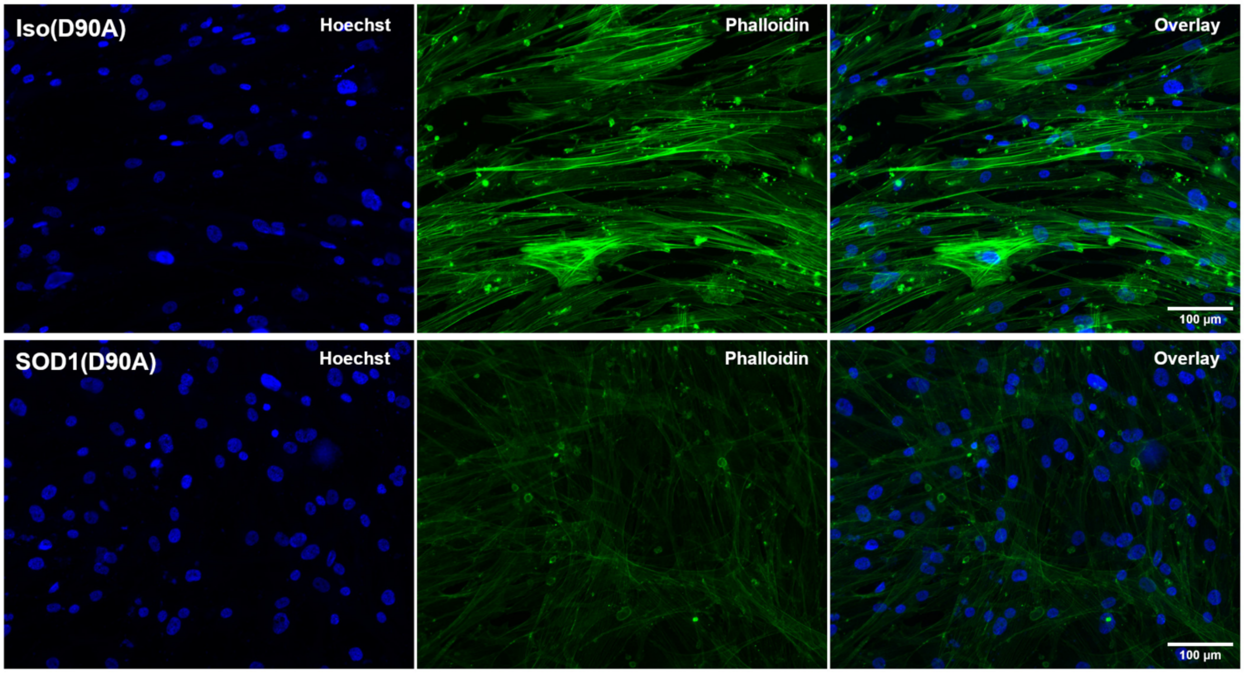


**Supplementary Figure 2.** Representative images of the actin cytoskeleton of *SOD1*(D90A) and matching isogenic astrocytes Iso(D90A) visualized by phalloidin staining (green); Hoechst counterstaining (blue) is shown.

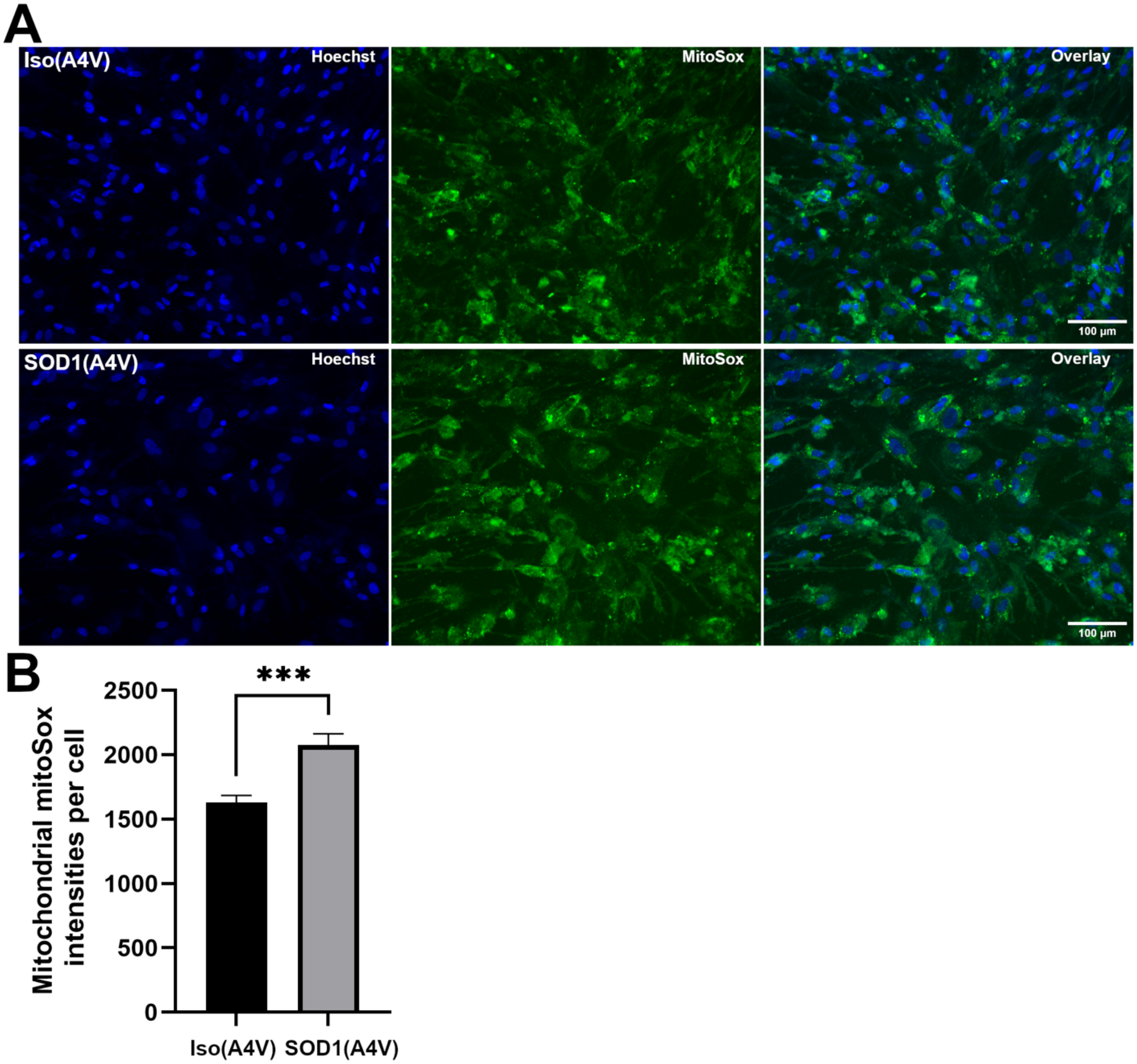


**Supplementary Figure 3. A)** Representative images of MitoSox (1 μM) fluorescence in *SOD1*(A4V) and isogenic astrocytes Iso(A4V), as indicated. **B)** Graph depicts quantifications of MitoSox intensities in *SOD1*-mutated astrocytes compared to matching isogenic astrocytes. Statistical analysis was performed with Student's t-test, graph shows mean ± *SEM*; ***$p < 0.0005$; $n = 3$ (more than 5,000 cells counted).
